# Study on the influence of G82S RAGE polymorphism on RAGE-Amyloid interaction in AD pathology

**DOI:** 10.1101/834374

**Authors:** Rani Cathrine. C, Bincy Lukose, P. Rani

## Abstract

Receptor for advanced glycation end products (RAGE) has been implicated in the pathophysiology of AD due to its ability to bind amyloid-beta and mediate inflammatory response. G82S RAGE polymorphism is associated with AD but the molecular mechanism for this association is not understood. Our previous *in silico* study indicated a higher binding affinity for mutated G82S RAGE, which could be caused due to changes in N linked glycosylation at residue N81. To confirm this hypothesis, in the present study molecular dynamics (MD) simulations were used to simulate the wild type (WT) and G82S glycosylated structures of RAGE to identify the global structural changes and to find the binding efficiency with Aβ42 peptide. Binding pocket analysis of the MD trajectory showed that cavity/binding pocket in mutant G82S glycosylated RAGE variants is more exposed and accessible to external ligands compared to WT RAGE, which can enhance the affinity of RAGE for Aβ. To validate the above concept, an *in vitro* binding study was carried using SHSY5Y cell line expressing recombinant WT and mutated RAGE variant individually to which HiLyte Fluor labeled Aβ42 was incubated at different concentrations. Saturated binding kinetics method was adopted to determine the K_d_ values for Aβ42 binding to RAGE. The K_d_ value for Aβ42-WT and Aβ42-mutant RAGE binding were 92±40 nM (95% CI-52 to 152nM; R^2^-0.92) and 45±20 nM (95% CI −29 to 64nM; R^2^-0.93), respectively. The K_d_ value of <100nM observed for both variants implicates RAGE as a high-affinity receptor for Aβ42 and mutant RAGE has higher affinity compared to WT. The alteration in binding affinity is responsible for activation of the inflammatory pathway as implicated by enhanced expression of TNFα and IL6 in mutant RAGE expressing cell line which gives a mechanistic view for the G82S RAGE association with AD.

## Introduction

Receptor for Advanced Glycation End-products (RAGE) belongs to the immunoglobulin superfamily, which interacts with various ligands and plays an important role in several pathological conditions [1]. Due to alternative splicing, various isoforms are generated such as full-length RAGE (fRAGE), secretory RAGE (sRAGE) and dominant negative RAGE (DNRAGE) and they bind to the ligands with similar affinity. The fRAGE consist of extracellular, hydrophobic transmembrane, and cytoplasmic domains, whereas sRAGE lacks a transmembrane domain. Extracellular domain has three immunoglobulins like domains namely variable (V) domain and two constant (C1 & C2) domains. The structural analysis of the ligand-binding domain within the V-domain structure of fRAGE indicates a hydrophobic cavity that is bordered by cationic residues and a flexible region (Thr55–Pro71). The flexible region allows further plasticity within the hydrophobic cavity, thereby promoting hydrostatic interactions with RAGE ligands [2,3]

Initiation of signal transduction upon the interaction of RAGE with its specific ligands helps in physiological processes such as chemotaxis, angiogenesis, inflammation, apoptosis, and proliferation [1,4]. The interaction of the same ligand with RAGE has different effects specific to the cell physiology where the activation of NF-kB helps in the survival of some cells and apoptosis of other cells [5]. As a multiligand receptor, fRAGE binds to the ligands like advanced glycosylation end products (AGEs), s100/calgranulins, amyloid-beta (Aβ) and amphoterin (HMGB1). Interaction of RAGE with AGEs results in acceleration of polymerization of Aβ, which increases the accumulation of insoluble plaques of Aβ, thereby enhances the risk of developing age-related disorders of CNS [6,7]. Excessive accumulation of these ligands tends to increase the inflammatory response and ROS production, resulting in cellular dysfunction.

RAGE, a potential contributor for neurodegeneration, has been implicated in accelerating degeneration and inflammation in neuronal tissues. The detrimental action of RAGE is exerted by its interaction with ligands which in turn activate the downstream pathways involving STAT, JKN, and NF-kB. Therefore, it has been indicated that the polymorphism within the ligand-binding domain of RAGE is associated with the activation of signal transduction pathways. There are several polymorphisms reported on the ligand-binding domain of RAGE. G82S polymorphism is one of the most frequently and naturally occurring single nucleotide polymorphisms (SNP) which enhances its affinity for ligand [8]. Thus, mutant expression shifts the signaling processes increases inflammation and contribute to several pathological conditions including Alzheimer’s disease (AD). Association of G82S RAGE polymorphism with AD is reported in Chinese [9,10], Korean [11] and the Turkish population [12]. Enhanced interaction between Aβ and fRAGE results in the activation of amyloid precursor protein (APP) cleaving enzyme that increases the production and the deposition of Aβ in the form of amyloid plaques [13]. Besides this, increased transport of circulating Aβ into the brain would be expected because RAGE has been shown to transport Aβ across the blood-brain barrier into the brain [14]. The structural determinants involved in post-translational modifications, such as N-glycans shown to affect RAGE binding and signaling, possibly by altering its association with various cell surface molecules. The V-type ligand domain of RAGE has two potential N-linked glycosylation sites (N25 and N81) and Srikrishna *et al.* [15] has demonstrated that N25 carries complex N-glycans while N81 may be unmodified or partially glycosylated with hybrid or high mannose glycans. G82S polymorphism also shown to affect glycosylation patterns in RAGE which could alter binding affinity to its ligands [16]. It is essential to understand the interaction of RAGE and Aβ, which would provide insight into its role in AD pathology and also to understand the molecular mechanism for the association of G82S RAGE polymorphism with AD.

The current study is designed to find RAGE-Aβ interaction scenario by comparing WT RAGE and G82S mutant RAGE through in silico and in vitro studies and to get a clear mechanistic view on the influence of glycosylation pattern on ligand binding affinity.

## Materials and methods

### Structures

X-ray crystallographic structure of monomeric RAGE ectodomain (PDB ID human 3cjj) is used in this study. G82S mutation was created and homology modeling of this RAGE variant was done using SCWRL4 with 3cjj as the template. All molecular structures were generated using Pymol. The glycans were virtually attached to the protein structure using the glycoprotein builder of GLYCAM web server (http://www.glycam.org). A complex glycan (Manα(1,6) [GlcNAcβ(1,2) Manα(1,3)]Manβ(1,4)-GlcNAcβ(1,4) [Fucα(1,6)]GlcNAcβ-OH) and a high mannose residue (Manα(1,2)-Manα(1,3)-Manα(1,6)-[Manα(1,3)]Manβ(1,4)-GlcNAcβ(1,4)-GlcNAcβ-OH) each having 7 carbohydrate units were added at the N25 and N81 position of the RAGE WT and mutant structures respectively.

### Molecular dynamics simulation

#### Force fields

MD Simulations were performed in Gromacs version 5.1. Glycans were modeled using the Glycam_06j-1 force field and the Amber ff12SB force field was used to model amino acid atoms. The resultant structures in amber topology were exported to gromacs topology using the modified version of ACPYPE.

#### Simulation setup

The structures of the glycosylated WT and mutant (G82S) form were simulated separately in a cubic periodic box with initial dimensions of 12.58 × 12.52 × 12.52 nm^3^ such that the minimum distance between each RAGE glycoform to the periodic boundary is 1.5 nm. Both the simulation system was solvated with an spc216 water Na+ or Cl^−^ ions, and then concentrated to 154 mM NaCl and energy minimized using steepest descent method. The energy minimized systems were equilibrated in the NVT ensemble followed by an NPT ensemble for 100 ps. The production dynamics were done in an isothermal-isobaric ensemble of 300K and 1 atm pressure for 50 ns. Both minimizations of energy were terminated using a maximum force tolerance of 1000 kJ mol ^−1^nm^−1^. The temperature, pressure, and NaCl concentration were chosen in such a way that mimics human body conditions. The resultant trajectory was analyzed using VMD and cavity volumes were tracked using MD pocket webserver with a grid spacing of 1Å

### RAGE gene amplification and purification

The peripheral blood mononuclear cells (PBMC) were isolated by HiSep LSM density gradient separation (Himedia, Mumbai, India) from human blood. Total RNA from PBMC was isolated using RNA-XPress™ reagent (Himedia, Mumbai, India). The quality of RNA was analyzed using nanospectrometer (Imple, USA). Total RNA (500ng) was then reverse-transcribed using RevertAid First Strand cDNA Synthesis Kit (Thermo Scientific, USA) and stored at −20° C until further use. The human fRAGE gene was amplified by using gene specific primers in 50μl reaction containing 100ng cDNA, 10pmol of each primer, and Ex-Taq polymerase (Takara Bio Inc., Japan).

Forward primer 5’TTA<u>GGTACC</u>ATGGCAGCCGGAACAGCAGT3’ and reverse primer 5’TAT<u>GAATTC</u>TCAAGGCC CTCCAGTACTAC 3’ were used for amplification of human RAGE gene. The underlined sequences represent *Kpn*I and *EcoR*I restriction sites, respectively. The expected product size is 1215 base pairs. PCR was performed and the product was purified from the 1% agarose gel using HiYield™ Gel/PCR DNA Mini Kit (Real Biotech Corporation, Taiwan).

### Cloning of WT RAGE gene and site-directed mutagenesis to create mutant RAGE variant

Cloning of the purified PCR product was done using InsTAclone PCR Cloning Kit (Thermo Fisher Scientific Inc., USA). The purified PCR product was ligated into pTZ57R/T vector and the recombinant vector was transformed into DH5α strain of *Escherichia coli.* From the subcultures, plasmid was isolated using GeneJET Plasmid Miniprep Kit (Thermo Fisher Scientific Inc., USA). DNA sequencing was performed in both directions using universal primers for M13 which flanks the multiple cloning site of the plasmid. Sequencing was outsourced to Eurofins Pvt. Ltd, Bangalore, India. The sequence-verified RAGE (WT) pTZ57R/T construct was used to perform mutagenesis. Site-directed mutagenesis (STM) was performed to create 82G to 82S change by mutagenic PCR using Q5 Site-Directed Mutagenesis Kit, New England Biolabs (NEB, England). Mutagenic primers (Forward primer 5’ CCTTCCCAACAGCTCCCTCTTC 3’, Reverse Primer 5’ ACACGAGCCACACTGTCC 3’) were designed using the tool available in http://nebasechanger.neb.com/. The steps involved in performing mutagenic PCR are as follows: Exponential amplification of recombinant clone, Kinases, Ligases and DpnI (KLD) treatment and transformation. The plasmid was isolated from transformed clones using the GeneJET Plasmid Isolation Kit (Thermo Scientific Inc, USA). Plasmid sequencing was outsourced to Bioserve Technologies Pvt. Ltd, Hyderabad to confirm the inserted sequence changes and also to ensure that no other mutations were created. The created G82S mutation was confirmed by restriction profiling of fRAGE gene PCR product using an enzyme AflIII and Alu I. The size of fRAGE gene is about ~1.2 kb and it was first digested with AflIII which cut at the position 433. The restricted product was electrophoresed on 2% agarose gel. Two bands corresponding to 433bp and 782bp was observed. 433bp sized band was eluted from the gel and digested with Alu I enzyme (5’ AGCT 3’). The digested product was electrophoresed on 4% agarose gel. When G is mutated to A at the position 250 of the fRAGE gene, this site was recognized by the Alu I enzyme which gives the size of 67bp, 182bpand184bp.

### Construct of expression cassette, cell culture, and transfection

The WT and mutant RAGE were excised from pTZ57R/T vector using *Kpn*I and *EcoR*I and cloned into pcDNA3.1 under the control of the CMV promoter. The transfection was performed in neuroblastoma cell line SHSY5Y procured from NCCS (Pune, India). Cells were cultured in modified Eagle’s medium (MEM, Himedia), supplemented with 10% fetal bovine serum (FBS, Gibco, Invitrogen) and antibiotics (streptomycin sulfate and benzylpenicillin) individually at final concentrations of 100 U/ml (Himedia, India). Cells were cultured at 37°C with 5% CO_2_ in tissue culture polystyrene dishes and transfected with pcDNA 3.1 recombinant vectors using jetPRIME kit, (Polyplus, France).

### Study on expression of recombinant RAGE variants SDS PAGE, Western blotting, and ELISA

Cells were lysed using RIPA lysis buffer (Himedia, Mumbai, India) and total protein concentration of cell lysate was determined by the BSA assay kit (Puregene, India). Cell lysate (15μg) were subjected to 12% SDS PAGE and western blotting. RAGE protein was detected using mouse monoclonal anti-RAGE antibody (E-1; Santa Cruz Biotech) and appropriate HRP-conjugated rabbit anti-mouse antibody at a dilution of 1:100 and 1:500 respectively. The beta-actin was detected using rabbit polyclonal anti-beta-actin antibody and secondary antibodies (HRP-conjugated Goat anti-rabbit antibody) were used at a dilution of 1:1000 and 1:500 respectively. RAGE protein in cell lysate was quantified using a commercially available ELISA kit from Quantikine R&D systems (DRG00, USA) according to the manufacturer’s instructions.

### Immunofluorescence

Cells were fixed in 4% paraformaldehyde and then permeabilized samples were blocked with 0.2% bovine serum albumin (BSA) for 45 minutes and incubated with primary anti RAGE antibody diluted in blocking buffer (0.2% BSA in PBS containing 0.02% Tween20, Santa Cruz Biotechnology, USA) for overnight at 4°C. Alex Fluor conjugated secondary antibody (Jackson Immuno research anti-mouse IgG) was added and incubated at 25°C for 45 min. The primary and secondary antibodies were used at a dilution of 1:200 and 1:100 respectively. Nuclei were counter-stained with DAPI for the determination of the viable cell number. The incubation medium was removed and the cells were stained for 10 min with DAPI (Invitrogen) in PBS with final concentration of 300nM. The specificity of binding was ascertained by performing the same procedure in the absence of the primary antibody (negative control). The samples were visualized using In cell 6000 microscope (GE Healthcare, USA). The images were background corrected using their corresponding negative controls and contrasted to the same levels using Image J software. The fluorescent images obtained were used to calculate the fluorescent intensity as shown in Fig 1 for the quantification of the expression of RAGE protein.

**Fig 1.**
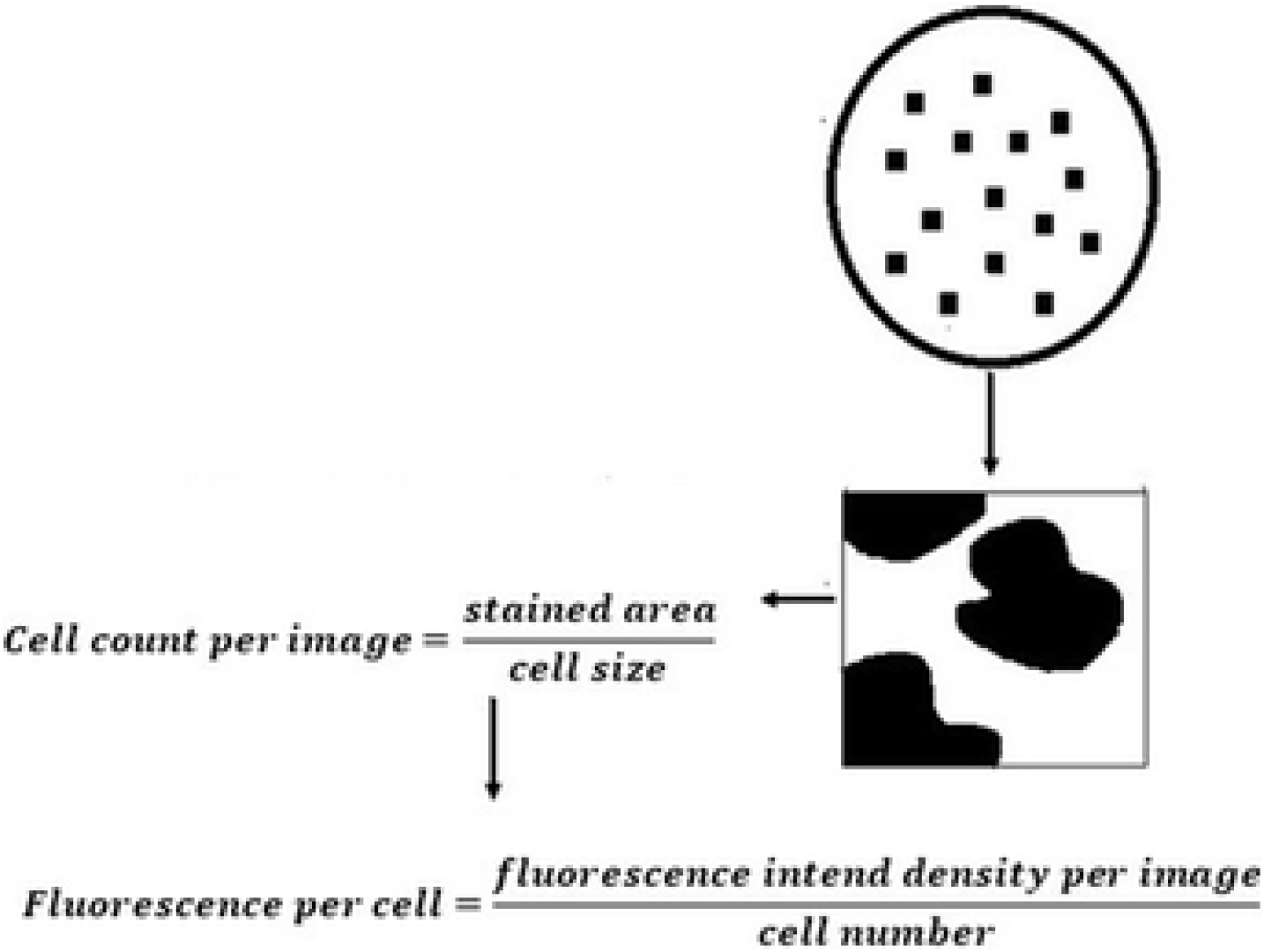
Calculation procedure followed to determine fluorescent intensity from fluorescent image obtained in cell analyzer.

### RAGE- Aβ interaction in transfected cell line

The experiment was performed to analyze the specific binding of Aβ to WT and mutant RAGE. We have used only Aβ42 since it is more pathogenic than other forms. To determine the optimal concentration for Aβ treatment, recombinant SHSY5Y cells were incubated with varying concentrations of Aβ42 (HiLyte Fluor labeled Aβ42, Anaspec) (50, 100, 150, 250, 500, 1000, 1250, 1500nM) for 3 h and washed to remove unbound ligands. The amount of Aβ42 bound to the cells was measured as mean fluorescent intensity using In Cell 6000 microscopy (GE healthcare, USA). The specific binding of Aβ42 was calculated by subtracting the fluorescence of cells in the absence of Aβ42 from that of fluorescence in the presence of varying concentrations of Aβ42. The data were analyzed using Image J software as described in Priesnitz *et al*. [17].

### Quantification of RAGE variants and inflammatory markers

To study the RAGE expression and inflammatory pathway activation due to RAGE-Aβ interaction, WT, and mutant RAGE transfected cells (untreated) and ligand treated cells were taken for the study. RAGE expression and inflammatory pathway activation was assessed by qPCR. The experiment was performed in single ligand concentration and concentration for treatment was decided from the saturation curve obtained from the binding study. The cells were treated with 1μM concentration of Aβ ligand for 3hrs and washed to remove unbound ligands. The cells were taken for the study after 48h and the experiments were performed in duplicate. Total RNA was isolated from transfected recombinant cells and ligand treated cells using RNA-XPress™ reagent (Himedia, Mumbai, India) according to the manufacturer’s instructions. Extracted RNAs were quantified by nanospectrometer, (Imple, USA). Total RNA (1μg) was reverse-transcribed using RevertAid First Strand cDNA Synthesis Kit (Thermo Scientific, USA) and in a total volume of 25 μL according to the manufacturer’s instructions. Primers used for qPCR for quantification of total RAGE (fRAGE), sRAGE, TNFα, and IL6 were represented in Table 1. The Glyceraldehyde-3-phosphate dehydrogenase (GAPDH) was used for internal normalization. RT-qPCR reactions were conducted in a 96-well plate using CFX96 Touch™ Real-Time PCR - Bio-Rad. Each reaction was performed in triplicate in 10 μL volume containing 1X SYBR Premix Ex Taq II (Takara Biotechnology), 50nM of each primer and 100ng of cDNA. The cycling conditions were as follows: 95°C for 10 sec, followed by 40 cycles at 95°C for 5 sec and 60°C for 30 sec. The transcript copy number of genes was determined based on their Ct values and the expression levels were calculated using 2^−ΔΔCT^.

**Table 1.**
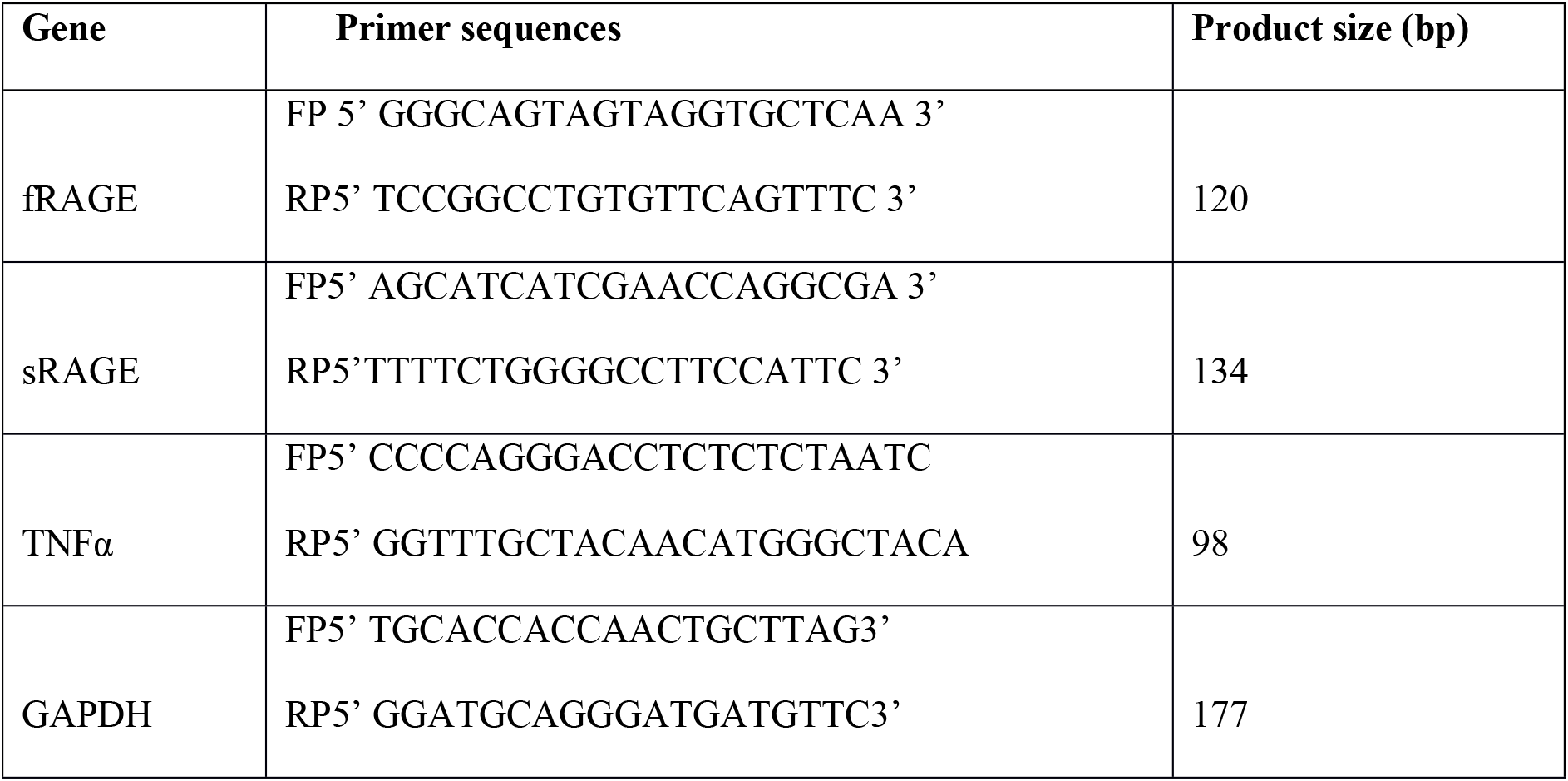
Primers sequence used for analysis of RAGE variants by qPCR.

## Results and discussion

### Conformational stability of G82S RAGE variant

The side-chain conformation prediction of the mutated structure (G82S) was done in SCWRL which is based on the minimization of the total energy of the entire model and the corresponding assignment of rotamers. For predicting the stability change caused by single point mutation STRUM was used. STRUM prediction is based on the difference in the free energy gap between wild type (ΔGm) and mutant protein (ΔGw), ΔΔG = ΔGm −ΔGw. A ΔΔG below zero indicates that the mutation causes destabilization [18]. ΔΔG value is negative for G82S RAGE (Table 2), which implies that G82S causes the destabilization of protein. Xie *et al*. [19] also reported that RAGE produced in bacteria lacking *N*-linked glycosylation due to G82S polymorphism causes a local change around the mutation site and a more global destabilization of the protein structure, with increased flexibility of the V-domain as shown by NMR spectroscopy.

**Table 2.** ΔΔG results from STRUM analysis.

| Protein | Polymorphic Position | Amino acid (wild type) | Amino acid (mutant type) | $\Delta\Delta G$ (Kcal/mol) |
| --- | --- | --- | --- | --- |
| RAGE | 82 | G | S | -1.99 |

Molecular dynamics has been carried out for 50 ns for both glycosylated form of WT and mutated RAGE. The initial and final conformations for both are shown in Fig 2A. When *t* = 0 ns, the conformation of both glycosylated WT and mutated RAGE is approximately the same and as time passes both the structures are deviating from the initial conformation. Root mean square deviation (RMSD) of the backbone atoms of the ectodomain of both wild type and mutated glycosylated RAGE structures are represented in Fig 2B. RMSD of glycosylated WT RAGE displays more fluctuations indicating that the conformational sampling of core backbone residues has not converged towards an equilibrium state within the 50 ns simulation period and has a lot of flexible side chain. However mutated RAGE variant attained stable conformation within the simulation time scale as evidenced by RMSD (Fig 2B). The secondary structure analysis by DSSP an inbuilt tool in GROMACS shown that most of the residues remained in β-sheet conformation throughout the simulation for both glycosylated systems (Fig 2C).

**Fig 2.**
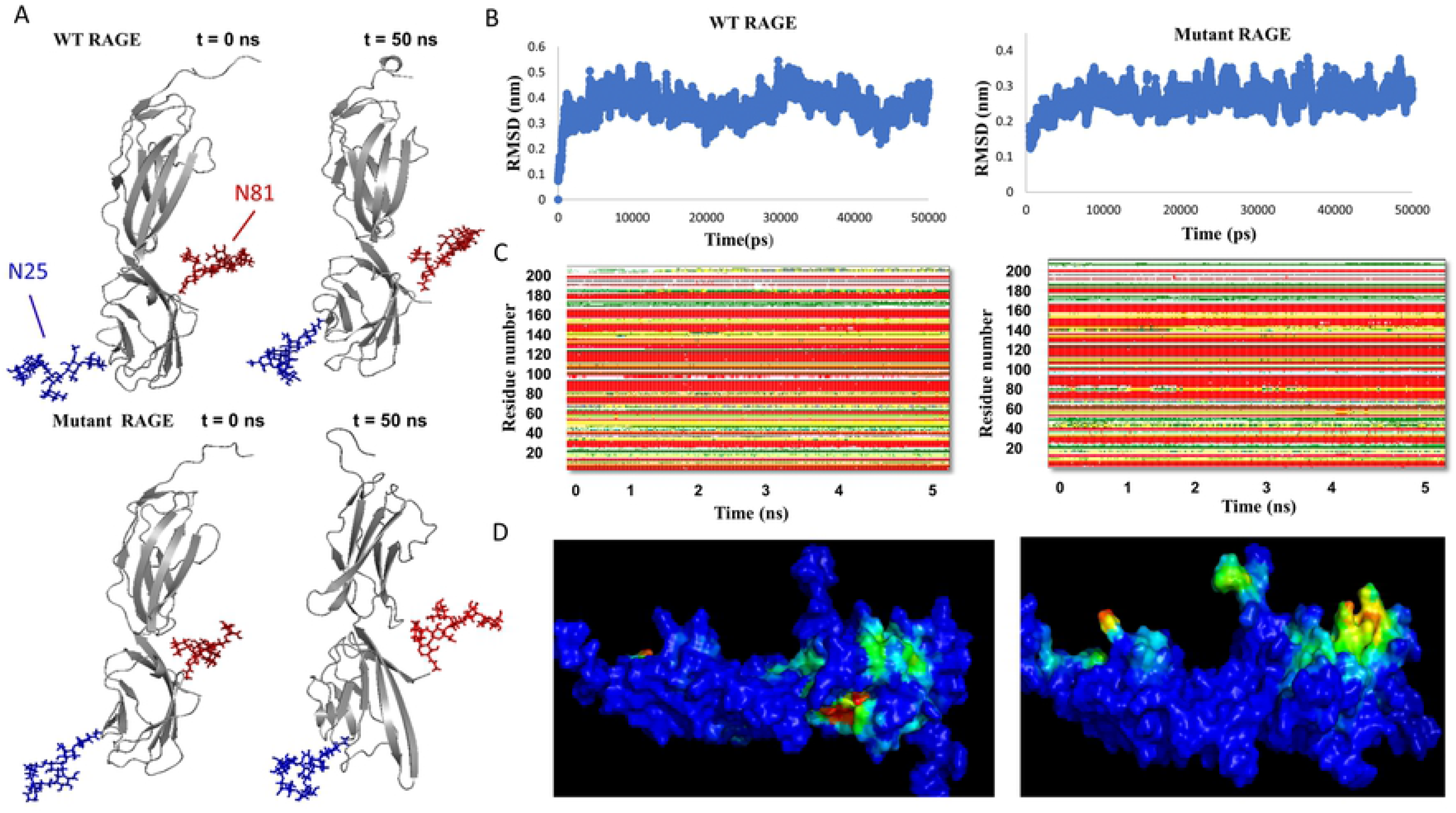
A) Initial and final conformations of the simulated glycosylated - WT RAGE, Mutant RAGE at t= 0 ns and t= 50 ns. B) Core backbone RMSD. Core backbone residues RMSD over 50 ns from the initial conformations for simulation of WT RAGE and mutant RAGE. C) DSSP analysis of glycosylated WT RAGE and mutant RAGE 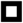 Coil, 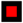 β-Sheet, 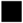 β-Bridge, 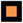 Bend, 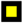 Turn, 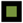 3-Helix. D) Binding pockets in glycosylated WT RAGE and mutant RAGE. Colors range from blue (low density = no particular cavity) to red (high density = conserved cavity).

### Binding pocket analysis

For tracking ligand/small molecule binding sites on the RAGE structures, MD pocket has been used which is based on the cavity detection algorithm. MD pocket detects transient sub pockets using an ensemble of crystal structures from molecular dynamics (MDs) trajectories. Cavity volumes were generated with an MD pocket using a grid spacing of 1 Å. It has been found that the cavity/binding pocket in the polymorphic variant of glycosylated RAGE (G82S) is more exposed /accessible to external ligands compared to WT RAGE which suggests that G82S polymorphism enhances the ligand-binding affinity of RAGE (Fig 2D).

In the present modeling study, G82S RAGE glycosylated at N81 and N25 showed a more exposed binding cavity compared to the glycosylated WT RAGE. The result gives preliminary evidence that anionic glycosylation at Asn81 may favor the electrostatic interactions with the cationic residues in the hydrophobic ligand-binding cavity that could be contributing to the flexibility in V domain thereby enhancing the affinity of G82S RAGE to Aβ peptides. Thus N-linked glycosylated Asn81 could play a major role in RAGE ligand binding, controlling the access and binding of ligand to the hydrophobic cavity.

### Site-directed mutagenesis of G82S RAGE gene confirmation

Genome-wide association study is widely conducted to help in developing a more accurate therapeutic and diagnostic target for various kinds of human diseases. Several epidemiological studies have been performed showing the association between RAGE polymorphism and various diseases namely rheumatoid arthritis [20], type 1 and 2 diabetes [21–23] and coronary artery diseases [24]. Functional SNP in RAGE namely G82S polymorphism is shown to be associated with increased risk for AD [9,10].

The gradient PCR with gene specific primers was used for amplification of fRAGE gene. The PCR product was electrophoresed in 1% agarose gel. Expected product size of 1215bp corresponding to fRAGE was observed in annealing temperature of 69 −72°C (Fig 3A). The purified PCR product was cloned into pTZ57R/T vector and sequence verified. The fRAGE gene was further cloned into mammalian expression vector - pcDNA3.1 (Fig 3B) and the cloned product was restricted to confirm product insertion (Figs 3 C and D) and the orientation of the cloned gene as shown in Figs 3 E and F.

**Fig 3.**
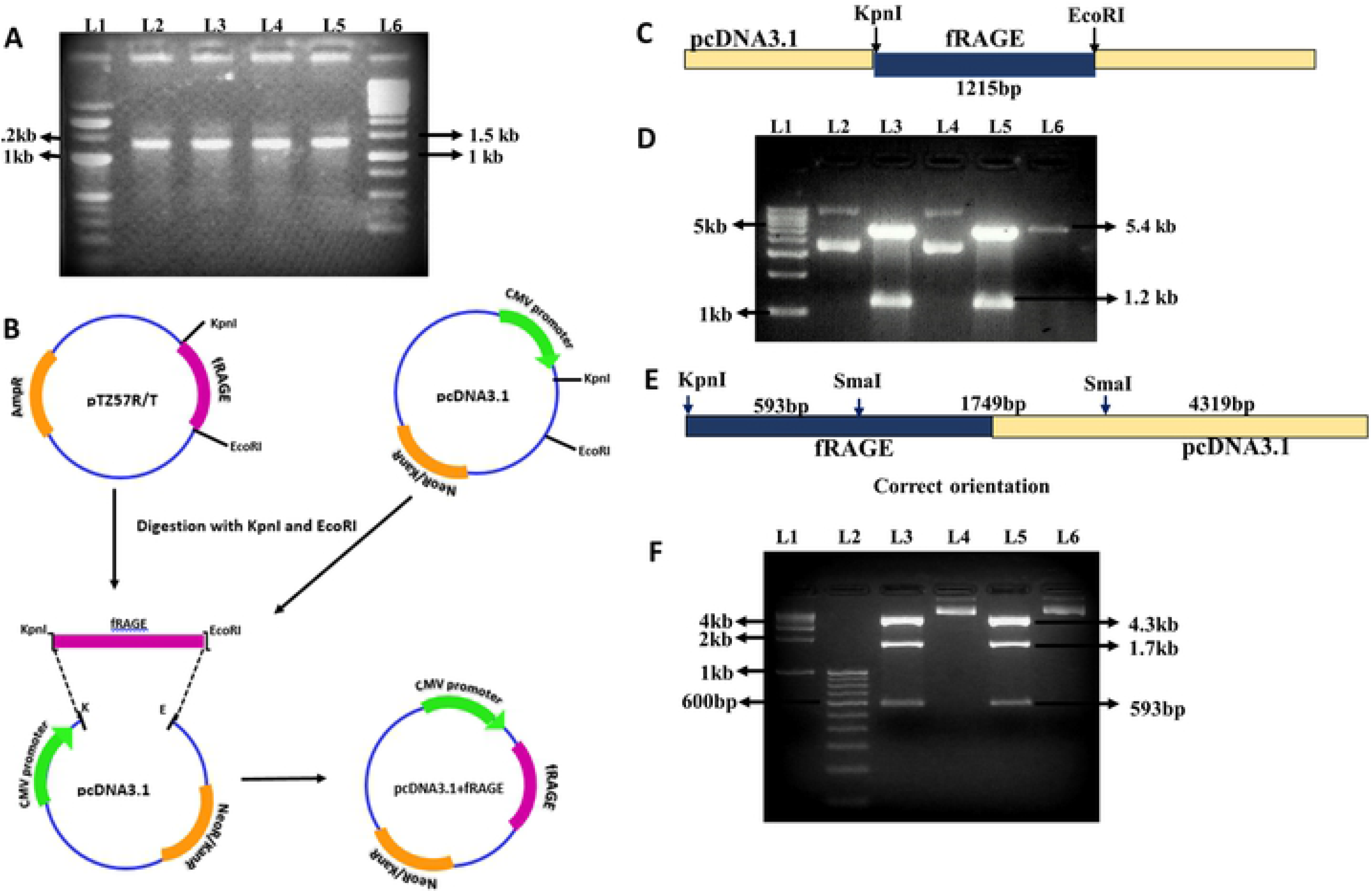
A) PCR product of amplified fRAGE gene. L1 −100bp marker; Lane 2-5 annealing temperature 69-72°C; L6 −1Kb marker. B) Schematic representation of recombinant pcDNA3.1 vector with fRAGE construct. The PCR product were cloned into pTZ57R/T vector. The pTZ57R/T fRAGE contract and pcDNA3.1vector was restriction digested using *Kpn*I and *EcoR*I and fragment was then inserted into pcDNA3.1 vector under the control of CMV promoter. C) Schematic representation of restriction pattern of recombinant pcDNA3.1 harbouring fRAGE gene. D) Restriction digestion of recombinant pcDNA3.1 construct with *Kpn*I and *EcoR*I were electrophoresed in a 1% agarose gel. L1-1kb DNA ladder, L2-pcDNA3.1 with WT RAGE (undigested), L3-pcDNA3.1 with WT RAGE (digested), L4-pcDNA3.1 with mutant RAGE (undigested), L5-pcDNA3.1 with mutant RAGE (digested), L6 - pcDNA3.1 vector (digested). Band size of 1.2kb shown in lane 3 & 5 indicates the presence of RAGE gene in recombinant pcDNA3.1 respectively. E) Schematic representation of orientation-based restriction pattern of recombinant RAGE gene in pcDNA3.1 vector. F) To confirm orientation of cloned RAGE gene recombinant pcDNA3.1 construct was restricted with *Kpn*I and *Sma*I and digested products were electrophoresed in a 2% agarose gel. L1-1kb DNA step ladder, L2-100bp DNA ladder, L3-recombinant WT RAGE pcDNA3.1 (digested), L4-recombinant WT RAGE pcDNA3.1 (undigested), L5-recombinant mutant RAGE pcDNA3.1 (digested), L6-recombinant mutant RAGE pcDNA3.1 (undigested). Band size of 593bp, 1750bp, 4272bp shown in lane 3 & 5 indicates the correct orientation of WT RAGE and mutant RAGE gene respectively in recombinant pcDNA3.1.

To create the G82S mutant RAGE variant, the site-directed mutagenesis was performed in pTZ57R/T with the WT RAGE construct (Fig 4A). The creation of the G82S RAGE variant was confirmed by restriction profiling and sequencing. The restriction digestion confirmed the substitution of A instead of G (Figs 4B, C, D and E). The sequencing results, which was analyzed using DNA baser assembler software. The switch from G to A in gene sequence and Glycine to serine in the translated sequence were confirmed as shown in Figs 4F, G and H.

**Fig 4.**
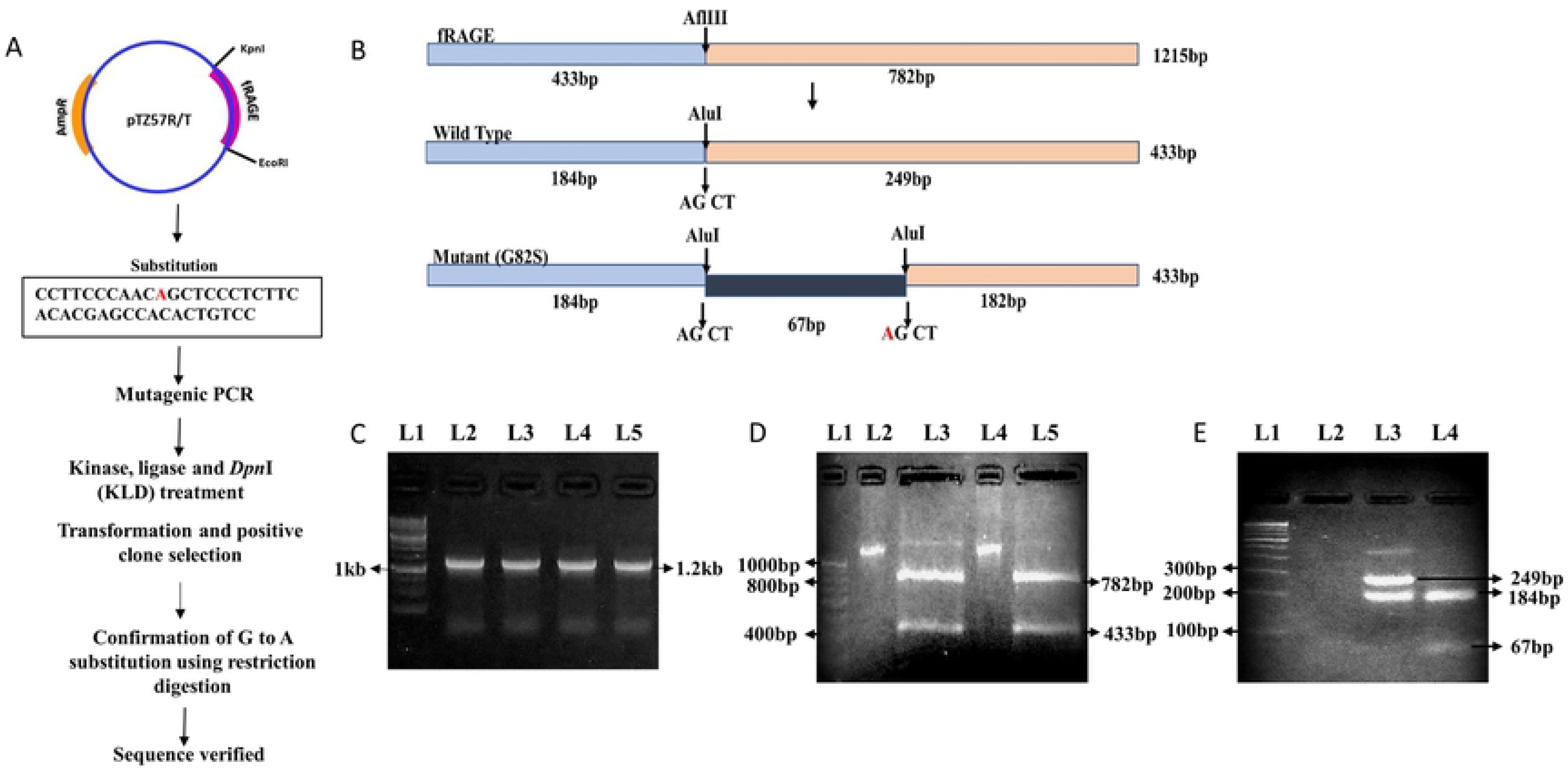
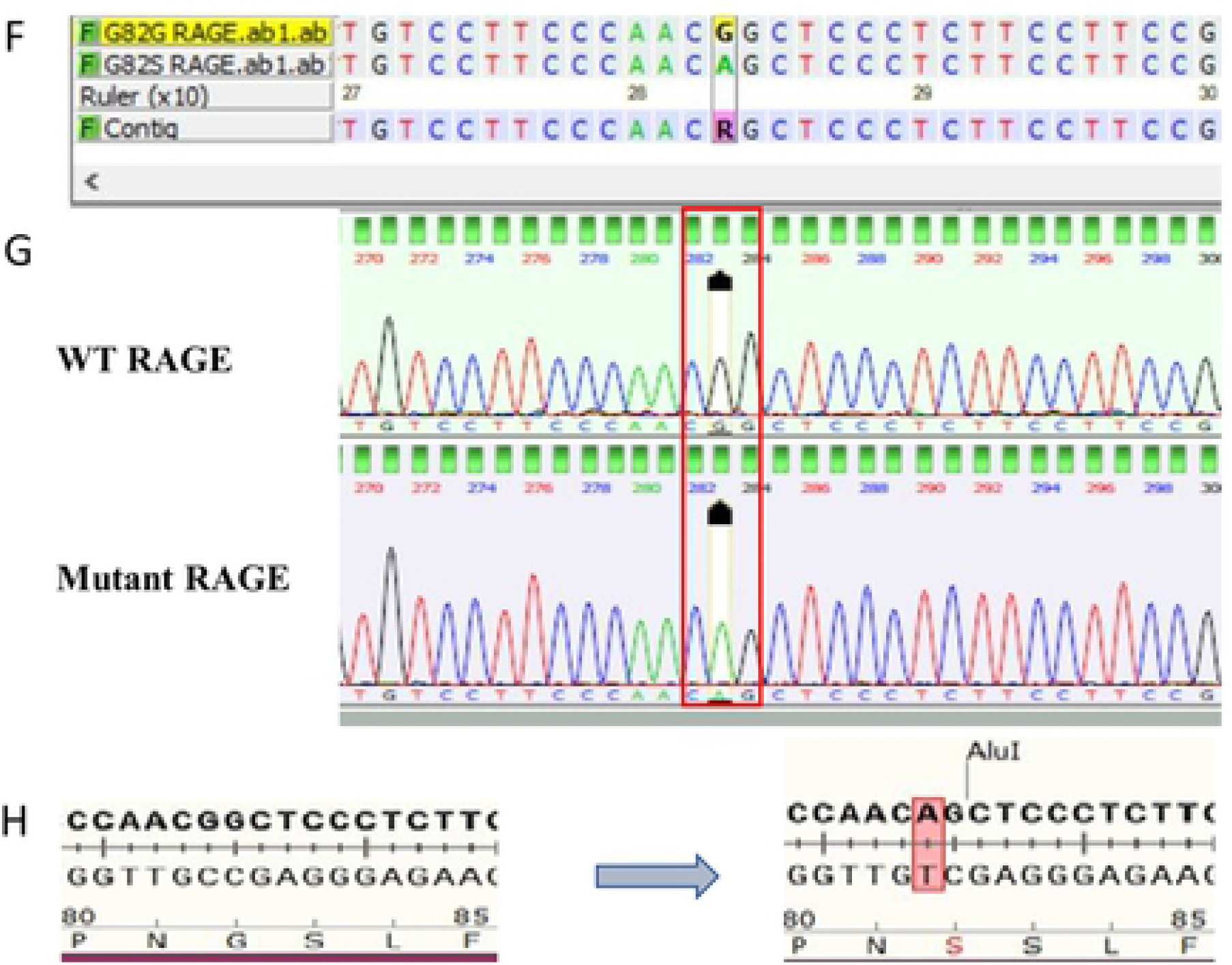
A) Schematic representation of site directed mutagenesis performed in pTZ57R/T RAGE construct. B) Restriction Mapping of WT & mutant RAGE with AflIII and Alu I enzyme. C) Amplification of WT & mutant RAGE gene. L1-1kb DNA ladder, L2 & 3 – WT RAGE PCR product, L4 & 5 – Mutant RAGE PCR product. D) Restriction profile of AflIII digested fRAGE gene PCR product. L1-100bp step up DNA ladder, L2-WT RAGE gene PCR product, L3-AflIII digested WT RAGE gene PCR product, L4-Mutant RAGE gene PCR product, L5-AflIII digested mutant RAGE gene PCR product. E) Restriction profiling of WT and mutant RAGE gene. L1-100bp step up DNA ladder, L2-empty, L3-Alu I digested WT RAGE gene fragment, L4-Alu I digested mutant RAGE gene fragment. F) Sequence confirmation of mutation in RAGE gene. Top row represents wild type nucleotide sequence and bottom row represent the mutant type sequence (G mutated to A in RAGE gene). G) Sequence chromatogram of WT and mutant RAGE sequence obtained from DNA baser assembler software. Red boxes highlight the sequencing location of the RAGE gene with G>A nucleotide change resulting in a G82S mutation. H) Representation of nucleotide change from G to A in RAGE gene which results in amino acid mutation at 82 position from Gly to Ser.

### Confirmation of RAGE expression in transfected SHSY5Y cell line

The WT and G82S mutant RAGE variants transfected and were expressed in transfected SHSY5Y cell line. The expression of RAGE was initially established by SDS-PAGE and western blotting of whole cell lysate (Figs 5A and B). The presence of 45kDA and 54KDa proteins indicates the expression of beta-actin and recombinant RAGE protein, which was confirmed through western blotting. The ELISA result of cell lysate also showed increased expression RAGE protein in transfected cells than non-transfected controls (Fig 5C). The mean fluorescent intensity was found to be 2-fold higher in transfected cell lines than empty vector-transfected cells and also non-transfected control cells (Figs 5D and E) confirming the expression of transfected RAGE gene.

**Fig 5.**
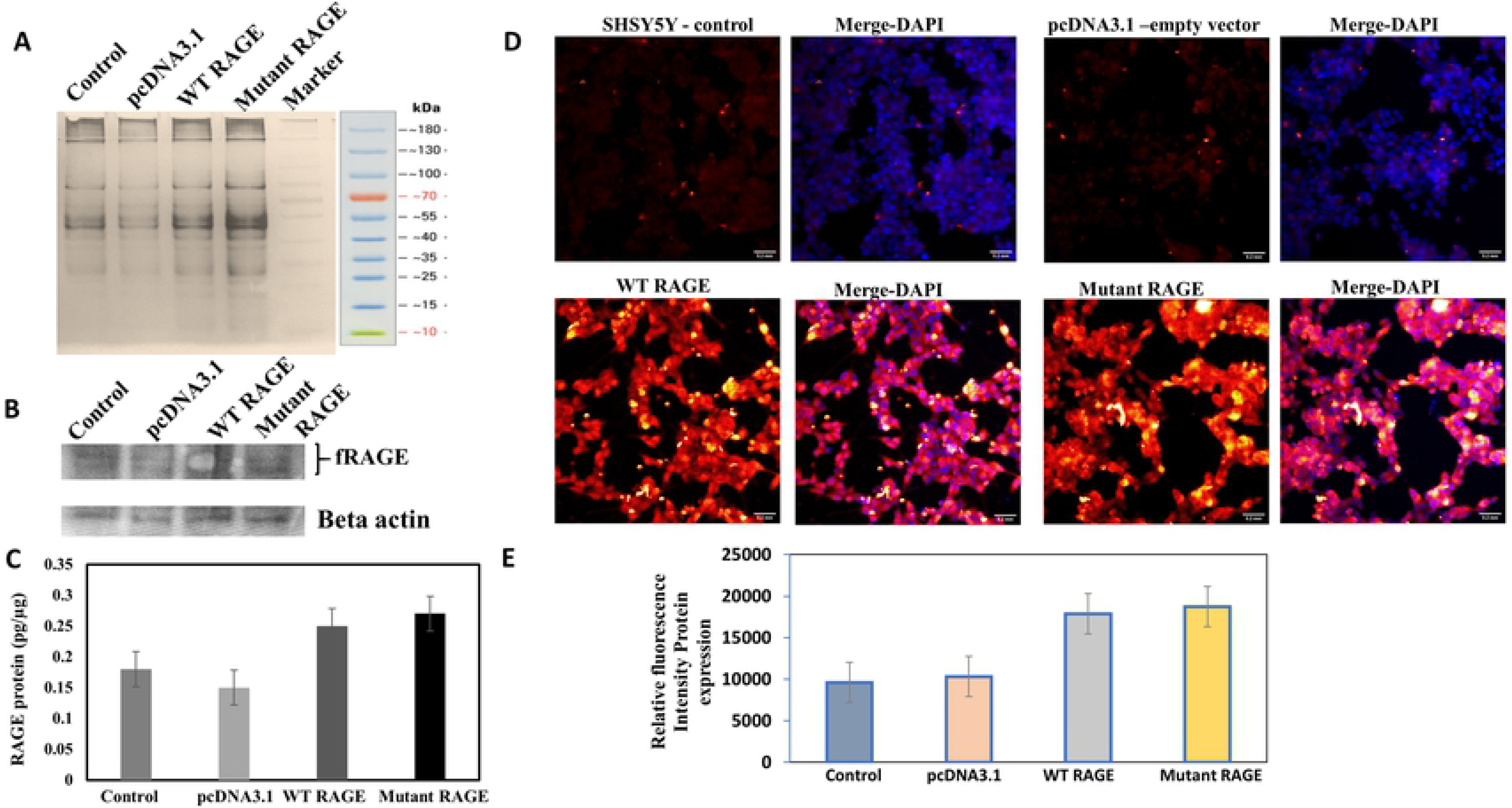
Protein profiling of transfected RAGE cell. A) SDS PAGE gel with cell lysate. B) Western blot analysis of RAGE in transfected and non-transfected cells. C) Quantification of RAGE expression using ELISA, bar graph depicts the mean RAGE expression level and error bars represents the standard error mean. D) Relative quantification of RAGE expression using fluorescent intensity obtained from fluorescent image of controls and transfected cells. E) Bar graph depicts the mean relative fluorescent intensity with error bars represents the standard error mean for RAGE protein expression. Cells were incubated with Alex Fluor tagged secondary antibody (30 min), DAPI (10min) and imaged. Scale bar correspond to 0.2mm in all images.

### G82S mutation in RAGE enhances Aβ42 binding affinity

Representative In Cell analyzer image of interaction of Aβ42 with RAGE in non-transfected, pcDNA3.1 transfected, recombinant WT and mutant RAGE transfected cells were given in Fig 6A. Expression of recombinant RAGE (WT and mutant) in SHSY5Y cells were confirmed through Alex Fluor tagged secondary antibody. The images of recombinant WT and mutant RAGE transfected cells incubated with varying concentrations of Aβ42 were represented in Figs 6B and C. which indicates a specific binding of Aβ42 to recombinant RAGE and also enhanced binding of Aβ42 to mutated RAGE. Binding of Aβ42 to RAGE increased up to 500nM concentration and saturated kinetics was observed beyond this concentration for both WT and mutant RAGE.

**Fig 6.**
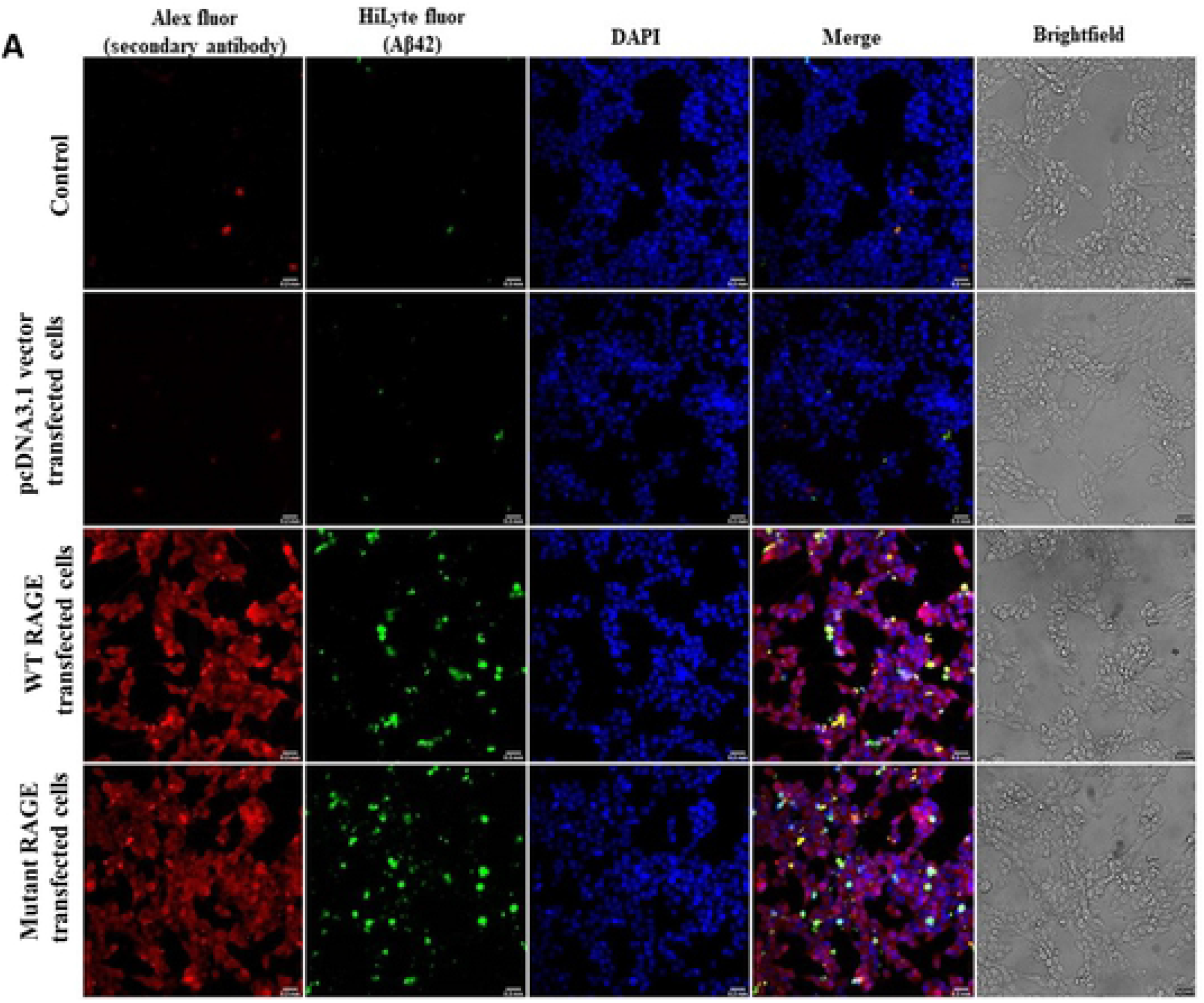
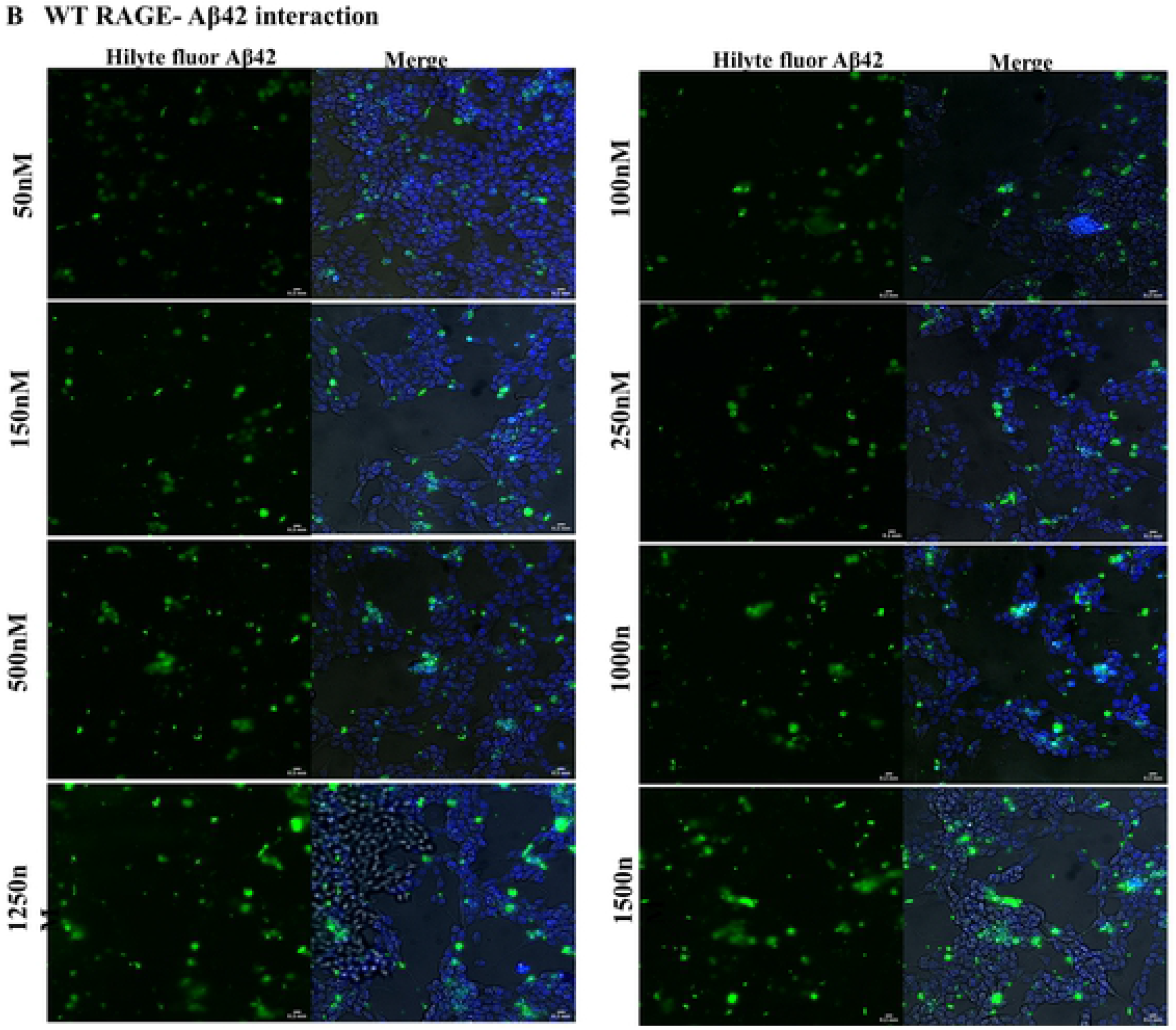
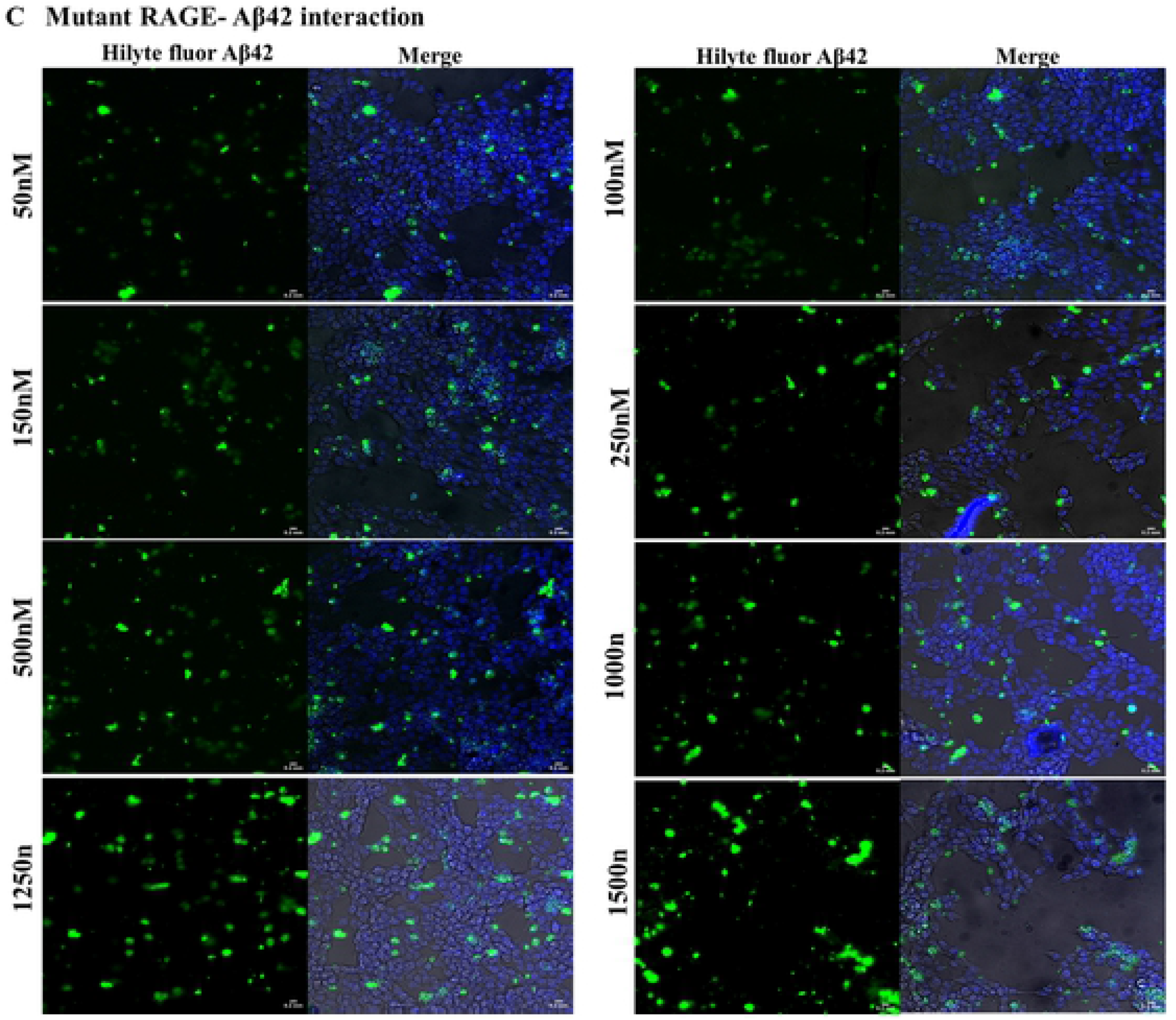
Interaction of Aβ42 with RAGE. A) representative image of controls and transfected cell lines immunostained with Alex fluor (secondary antibody) and DAPI after treating with HiLyte fluor labelled Aβ42 (1μM). B) Recombinant WT and C) mutant RAGE expressing SHSY5Y cells were incubated with varying concentrations of Aβ42 (50-1500 nM) (HiLyte Fluor labelled Aβ42, Anaspec).

A saturated binding kinetics method was adopted to determine the K_d_ value for Aβ42 binding to RAGE. The K_d_ value for WT RAGE and mutated RAGE were 92nM ±40nM (95% CI-52 to 152nM; R^2^-0.92) and 45nM±20nM (95% CI −29 to 64nM; R^2^-0.93; *p*<0.05) respectively which indicates that both RAGE variants are high-affinity receptor for Aβ42. K_d_ value for mutated RAGE was lower than WT RAGE indicating a significant increase in affinity for mutated RAGE for Aβ42 binding than WT RAGE (Figs 7 A, B and C). This explains the enhanced function associated with RAGE variants with G82S polymorphism.

**Fig 7.**
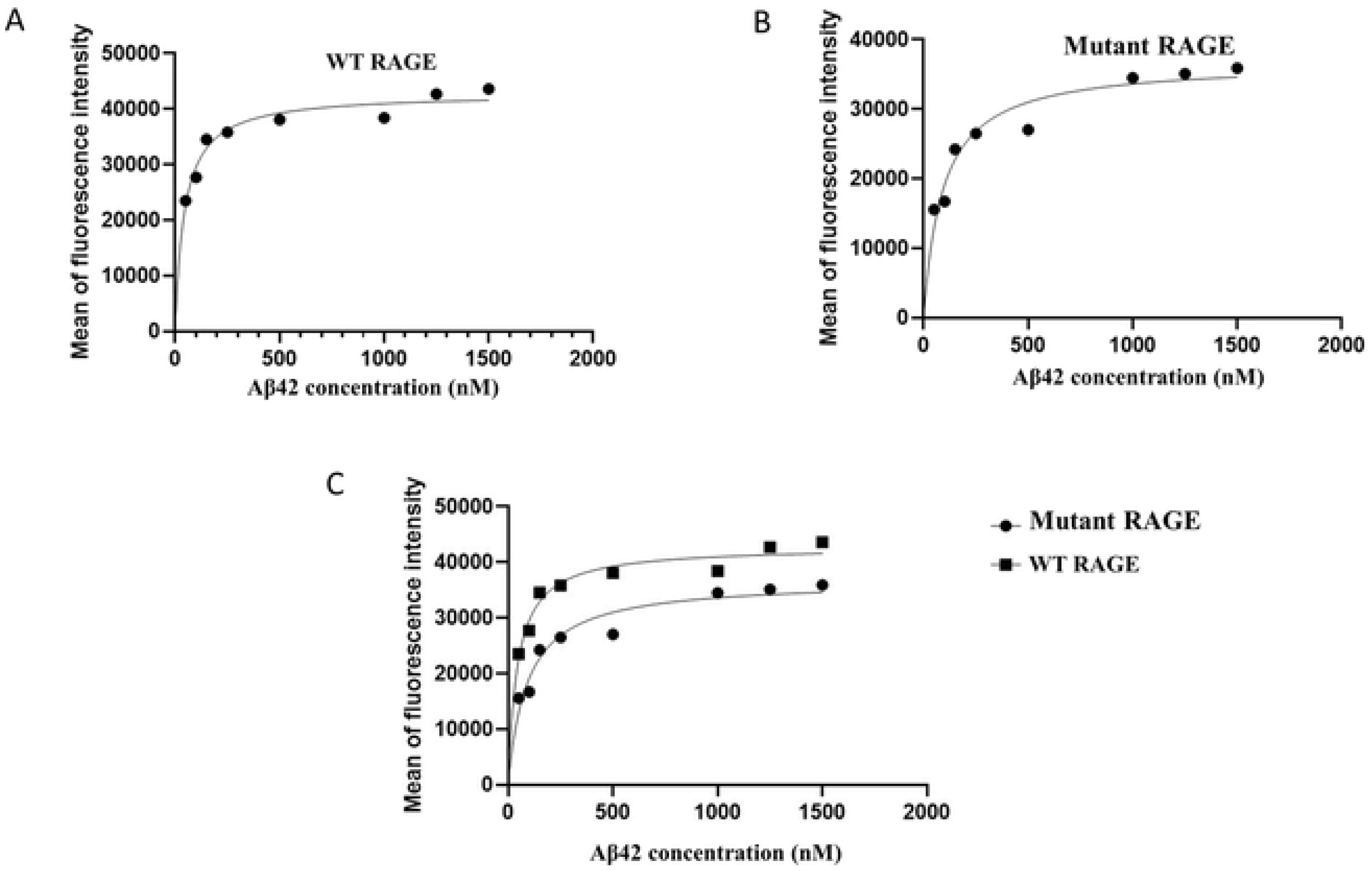
Binding curves of HiLyte Fluor – labelled Aβ42 interaction to WT RAGE and mutant RAGE in SHSY5Y cell lines. A) WT RAGE; B) mutant RAGE; C) Merge of WT and mutant RAGE. Transfected cells were incubated with increasing concentration of Aβ42 (HiLyte fluor labelled ligand). To calculate K_d_ of the interaction of Aβ42 to RAGE the mean fluorescent intensity of the Aβ42 bound vs. Aβ42 concentration added was fit to the equation Y=Bmax X/(K_d_ + X) using GraphPad prism software. The values are the average of five trails.

This G82S polymorphism occurs nearby to Asn81 one of the potential N-linked glycosylation sites. It is hypothesized that this modification plays a major role in RAGE-ligand interaction. This might be because glycine at 82^nd^ position is more flexible than serine. Also, it gives a probable clue that in WT RAGE, the N81th position may not be glycosylated. G82S polymorphism might stabilize the N-linked glycosylation at N81 thereby giving structural stability to the mutated RAGE than glycosylated WT RAGE. Previously, it has been shown that Asn25 in WT RAGE is always modified with fully processed N-linked glycan, whereas Asn81 is not favored for N-linked glycosylation. More recent studies indicate that Asn81 is also glycosylated in G82S polymorphic RAGE and this polymorphism might affect RAGE glycosylation [15].

### RAGE-Aβ interaction influences expression of RAGE variants

The NFκB activation is mediated by upstream pathways including RAGE-amyloid interaction and once NFκB is activated it creates an alteration in RAGE expression. To study this alteration in the expression of RAGE isoforms, qPCR was performed to quantify fRAGE and sRAGE. The qPCR results showed a similar range of fRAGE expression in both WT and mutant RAGE expressing cells when not treated with Aβ42, whereas the expression levels in Aβ42 treated cells showed a marginal increase in expression (Fig 8A). The enhanced expression of fRAGE could be a result of a positive loop mechanism of upregulating the expression of RAGE by amplifying the cellular response due to external stress. This notion is supported by the finding that fRAGE expression is increased in AD brains [25,26].

**Fig 8.**
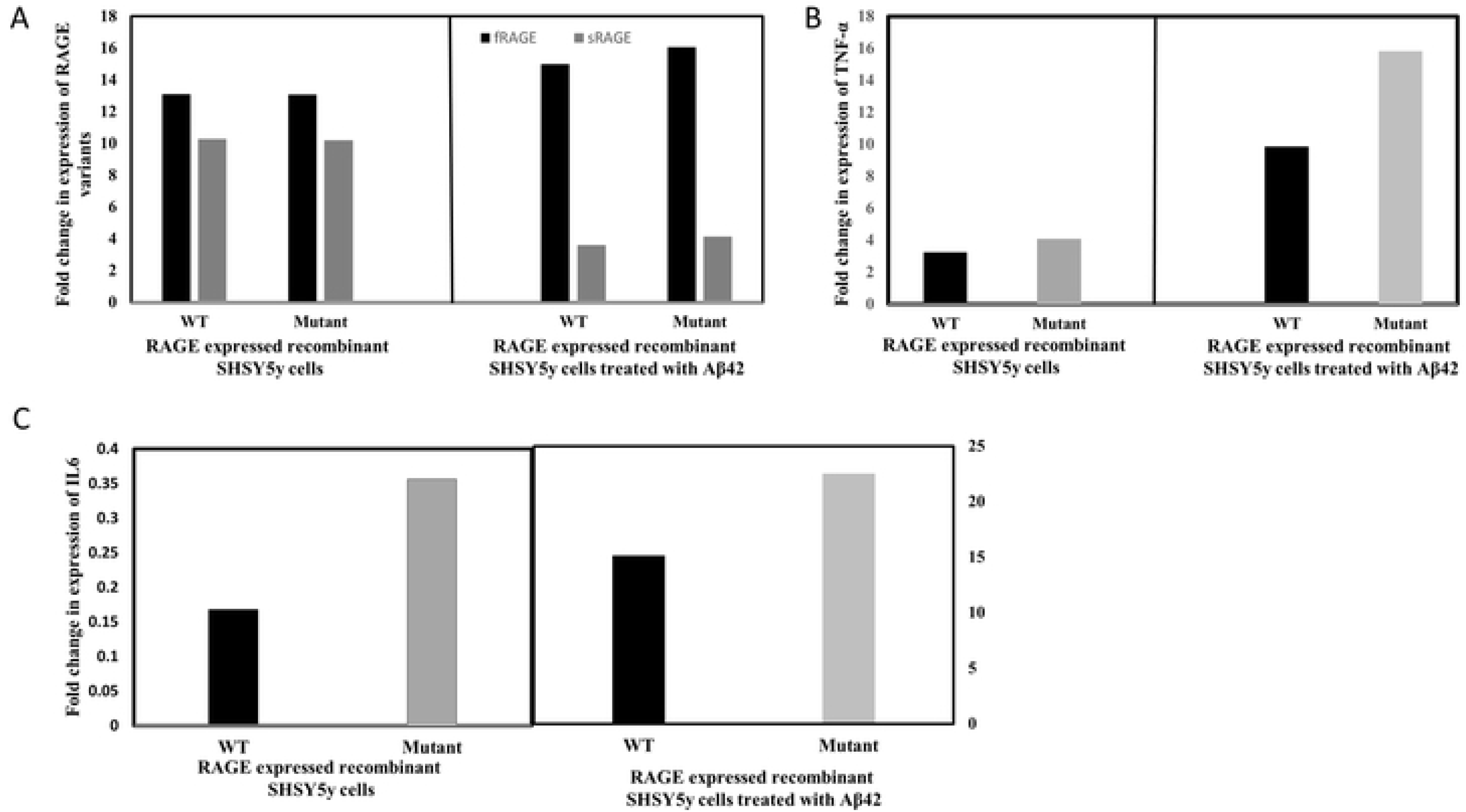
Relative quantification of recombinant RAGE variants and Inflammatory markers expression using qPCR. A) RAGE variant expression; B) TNF-α; C) IL6. Bar graph depicts the mean gene expression levels and expression values are obtained by 2^−ΔΔCT^.

sRAGE expression is higher in ligand untreated cells which decreased drastically upon Aβ42 treatment. The decreased expression of sRAGE could be due to alteration in the expression of RAGE isoforms. Besides, it's been shown that 82S carriers have roughly half the maximum amount of sRAGE when compared with 82G carriers [9, 11], implying that the increased ligand affinity of RAGE receptor leads to a dysregulation of RAGE isoforms. Since sRAGE acts as a decoy receptor for Aβ binding and decrease in sRAGE expression during Aβ treatment might decrease Aβ clearance and further lead to Aβ burden. This G82S polymorphism along with Aβ burden could thereby enhance fRAGE production, thus giving little room for sRAGE to exert its proposed protective mechanisms. Thereby both mechanisms namely decreased sRAGE expression and increased fRAGE expression can contribute to AD pathogenicity.

### RAGE-Aβ interaction elicits inflammatory response

The RAGE receptor which binds to a variety of proinflammatory ligands transmits the signal from the ligand to NFκB regulated cytokines production. To confirm the inflammatory pathway activation due to RAGE-Aβ interaction, the cells were exposed to Aβ42 and tested for cytokines levels (TNFα and IL6). The qPCR results confirmed the increased expression of pro-inflammatory cytokines upon Aβ42 interaction. Comparatively higher expression of TNF-α (9.8-fold), IL6 (15.13-fold) were observed in Aβ42 treated cells than untreated cells. A similar trend was also observed in the mutant RAGE expressed cell line. The expression of TNFα and IL6 were higher in mutated RAGE expressed cells than WT RAGE as shown in Figs 8B and C.

The RAGE-Aβ interaction induces the inflammatory pathways as demonstrated by an increased expression of proinflammatory cytokines such as TNFα and IL6. The observation confirms that RAGE – Aβ interaction evokes a cascade of downstream pro-inflammatory signaling pathways and this effect is more prominent in G82S RAGE polymorphism.

## Conclusions

In our study, we report that G82S polymorphism stabilizes RAGE glycosylation at Asn81 suggesting that the increase in flexibility of the V-domain caused by the global destabilization effect of G82S mutation might be the cause for more exposed binding cavity in polymorphic glycosylated RAGE which enhances the ligand-binding affinity of RAGE towards Aβ.

Our study suggests that expression of RAGE is increased at sites of Aβ42 accumulation and polymorphisms within ligand-binding regions (G82S) alters RAGE variant expressions leading to enhanced fRAGE and decreased sRAGE expression thereby amplifying the inflammatory response. These results cumulatively suggest that RAGE is a potential candidate for a therapeutic approach in AD. This can be envisaged by using sRAGE therapeutically to clear Aβ or by using antibody complimentary to Aβ binding region of fRAGE to prevent inflammatory process in AD.

